# Improving biome labeling for tens of thousands of inaccurately annotated microbial community samples based on neural network and transfer learning

**DOI:** 10.1101/2022.09.09.507244

**Authors:** Nan Wang, Teng Wang, Kang Ning

## Abstract

Microbiome samples are accumulating at a fast speed, leading to millions of accessible microbiome samples in the public databases. However, due to the lack of strict meta-data standard for data submission and other reasons, there is currently a non-neglectable proportion of microbiome samples in the public database that have no annotations about where these samples were collected, how they were processed and sequenced, etc., among which the missing information about collection niches (biome) is one of the most prominent. The lack of sample biome information has created a bottleneck for mining of the microbiome data, making it difficult in applications such as sample source tracking and biomarker discovery. Here we have designed Meta-Sorter, a neural network and transfer learning enabled AI method for improving the biome labeling of thousands of microbial community samples without detailed biome information. Results have shown that out of 16,507 samples that have no detailed biome annotations, 96.65% could be correctly classified, largely solving the missing biome labeling problem. Interestingly, we succeeded in classify 250 samples, which were sampled from benthic and water column but vaguely labeled as “Marine” in MGnify, in more details and with high fidelity. What’s more, many of successfully predicted sample labels were from studies that involved human-environment interactions, for which we could also clearly differentiated samples from environment or human. Taken together, we have improved the completeness of biome label information for thousands of microbial community samples, facilitating sample classification and knowledge discovery from millions of microbiome samples.

## Introduction

With the development of next generation sequencing technology, the annual sequencing data has grown rapidly and the data volume is huge. Utilizing Metagenomics Sequencing and 16S sequencing and annotation technology, we could obtain the taxonomic structure and abundance information about microbial communities (Escobar-Zepeda et al. 2015). The analyses of these microbial data play a very important role in environmental protection (Bahram et al. 2018), water pollution monitoring (Wang et al. 2020), and disease diagnosis and prevention (Halfvarson et al. 2017; Gaulke and Sharpton 2018), etc. For example, studies on indoor microbial communities have provided insights on how human-environment interactions could affect the taxonomic structure patterns of households (Kelley and Gilbert 2013), as well as newly opened hospitals (Lax et al. 2017). Currently, paramount microbiome samples have been deposited to multiple public databases (Keegan et al. 2016; Gonzalez et al. 2018; Mitchell et al. 2020; Sayers et al. 2021), out of which EBI-MGnify (or MGnify) offers an automated pipeline for the analysis and archiving of microbiome data to help determine the taxonomic diversity and metabolic potential of environmental samples. The millions of microbiome samples deposited in these public databases could facilitate the comparison, clustering and mining of the microbiome data.

The ideal microbiome database is one in which all datasets are annotated exhaustively, clearly and precisely. However, due to the absence of a strict and unified submission standard, as well as the complex sources of samples at the outset of the database construction, a considerable number of sample annotations are “rough sketches” with imprecise or non-detailed source labels. For instance, a considerable proportion of samples which should’ve been attributed to diverse microbial categories were not annotated in detail, but were classified as “Mixed biome”. The rough sketches are primarily comprised of three types of improperly annotated samples: un-annotated samples (samples annotated as “Mixed biome”), under-annotated samples (samples with coarse annotations that could be refined) and mis-annotated samples (samples with incorrect annotations). Since the establishment of the database, the total sample size of MGnify has increased significantly, and the proportion of inaccurately annotated samples, taking un-annotated samples for example, has also increased (**Fig. 1**), which can trigger severe problems listed as follows.

**Figure 1.**
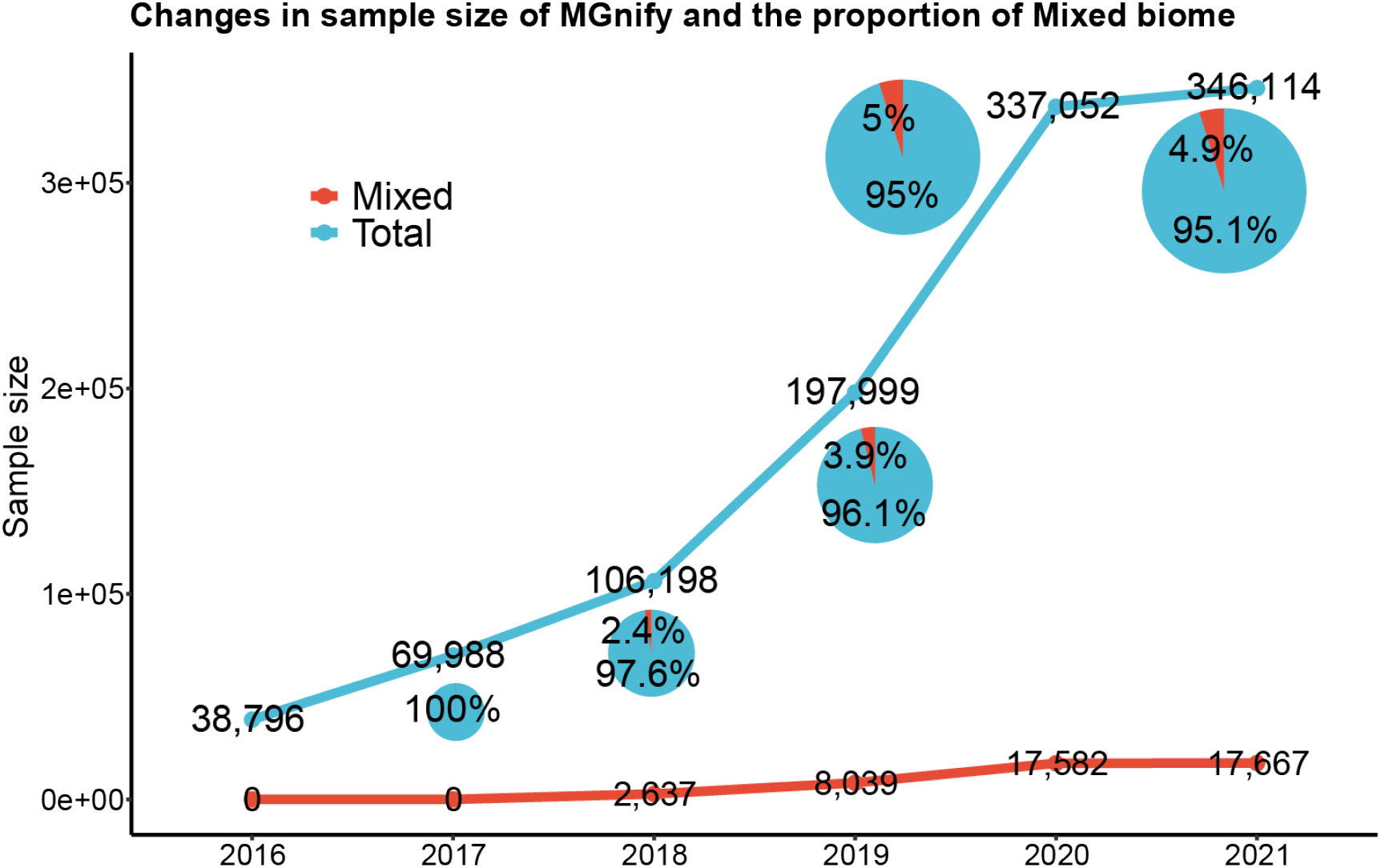
The increasing number of all samples and those annotated as “Mixed biome” in MGnify. The statistic of the annual amount of all samples and samples annotated as “Mixed biome” in MGnify from 2016 to 2021. The lines show the change of the annual amount of all sample and samples annotated as “Mixed biome”. The pie charts show the annual proportion of the samples annotated as “Mixed biome” in all samples.

Firstly, inaccurate annotations could lead to a substantial proportion of Microbiome samples being wasted. The increasing number of samples annotated as “Mixed biome” have a high potential to be wasted due to the inconvenient mining of these public datasets. For example, due to the annotations of “Mixed biome,” unannotated infant gut samples have a high probability of being completely ignored, which hinders the examination of the dynamics of human gut microbial development (study accession: MGYS00005507) (Brooks et al. 2014). Secondly, inaccurate annotation will result in the misapplication of microbiome samples or even the failure of the research. Due to the lack of strict meta-data standard for data submission, there are currently a massive part of under-annotated samples existing in the databases. For example, if animal gut flora related samples are coarsely annotated as host-associated samples, these samples may be incorrectly adopted as human gut flora related samples when conducting the data mining phase of a disease associated study. In the absence of sufficiently effective means of data cleaning, these data can end up being misused and therefore hinder the drawing of valid research conclusions or even lead to study failure. Thirdly, inaccurate annotations can lead to “cascading accumulation” of errors when included in the secondary databases. Recently, numerous efficient secondary or specialized databases have emerged, such as GM-repo (Wu et al. 2020), linking data from different primary databases. Inaccurate annotated samples exist in different primary databases, once entered into secondary databases or even re-annotated, the true origin of the samples will be unverifiable and further increase the number of inaccurately annotated samples in secondary databases, leading to “cascading accumulation” of errors. With the ever-increasing proportion of samples accumulated in the current microbiome databases (**Fig. 1**), the aforementioned issues pose a significant obstacle for microbiome research. Therefore, highly intelligent and automatic method that could disentangle and refine the biome information for microbiome samples is desirable.

In this study, we have proposed Meta-Sorter, which was designed based on neural network (Kriegeskorte and Golan 2019) and transfer learning (Cai et al. 2020), to disentangle the biome labels for samples with inaccurate biome annotations. A neural network model was first constructed based on 94,874 samples introduced into MGnify before January 2020 (existing samples) with detailed biome annotations, and showed high robustness and accuracy (average AUROC=0.896). This model was then applied on all existing samples annotated as “Mixed biome” for their detailed biome prediction, with results showing that 95.41% samples were consistent with their meta-data. Secondly, due to the reduced accuracy of the neural network model on samples introduced into MGnify after January 2020 (newly introduced samples), we designed a transfer neural network model based on transfer learning, with the classification accuracy (average AUROC) again boosting to 0.989, and 97.62% of newly introduced samples annotated as “Mixed biome” correctly predicted. Combined the results on existing samples and newly introduced samples, we found that out of 16,507 samples that have no detailed biome annotations, 96.65% could be correctly classified, largely solving the missing biome labeling problem. Finally, we assessed practical application performance of Meta-Sorter on several concrete cases, such as differentiating the actual sources of samples only labeled as “Marine” in MGnify into benthic and water column, and classifying samples from studies that involved human-environment interactions into environment or human. Collectively, we have designed Meta-Sorter as a highly intelligent and automatic method that could disentangle and refine the biome information for microbial samples. Meta-Sorter is thus a useful tool for better classification and knowledge discovery from millions of microbiome samples.

## Results

### The workflow of Meta-Sorter

The aim of Meta-Sorter is to decode the biome labels (representing where the samples were collected) for samples without detailed biome information to provide more valuable information for researchers, by using neural network and transfer learning techniques. In this study, Meta-Sorter has two input files: a biome ontology (a hierarchical structure that represents the taxonomic hierarchy of the samples, for example, the biome “root-Host-associated-Human-Digestive system-Large intestine-Fecal” is represented in the ontology as: layer-1 represents “root”, layer-2 represents “Host-associated”, …, layer-6 represents “Fecal”), and the taxonomic structures (generally obtained by 16S sequencing or Metagenomics Sequencing and annotated by software such as QIIME (Bolyen et al. 2019) and KRAKEN (Lu and Salzberg 2020)) for microbial community samples with detailed biome information. These two input files are used to optimize the parameters of the neural network model by forward and backward algorithm. Inputting taxonomic structures of microbial samples from unknown sources from the input layer, we could obtain the detailed source contribution of the samples from the output layer (**Fig. 2A**).

**Figure 2.**
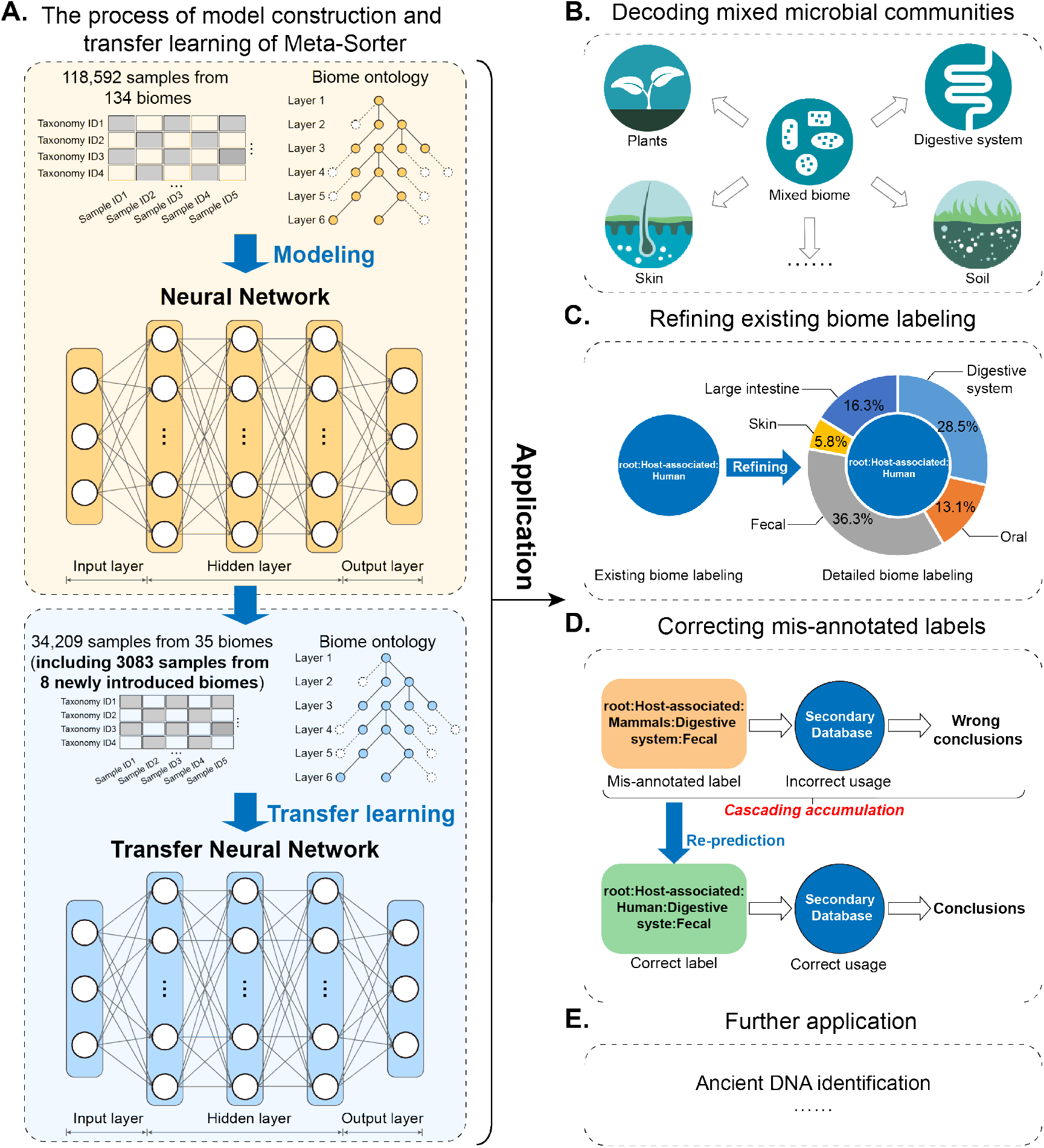
The rationale and applications of Meta-Sorter. (A) The process of model construction and transfer learning of Meta-Sorter. Two input files, which are biome ontology and samples’ taxonomic structures with detailed biome information, are required in the process of model construction and transfer learning. The yellow box shows that the neural network model was constructed based on 118,592 existing samples with detailed information of 134 biomes and their biome ontology. The blue box shows that the transfer neural network model was constructed by using 34,209 newly introduced samples from 35 biomes (including 3,083 samples from 8 newly introduced biomes) and transfer learning to the existing neural network model. (B-E) The applications of Meta-Sorter. (B) Meta-Sorter decoded the samples’ biome labels which were annotated as “Mixed biome” into detailed biome labels. (C) Meta-sorter refined the biome labels more detailed to obtain more valuable information for reference. (D) Meta-Sorter corrected the mis-annotated samples’ labels to avoid “cascading accumulation”. (E) Further more application of Meta-Sorter.

Meta-Sorter has a neural network model constructed based on 118,592 microbial samples from 134 biomes and their biome ontology. Notice that in this study, the 118,592 samples with detailed biome information used here were those deposited into the MGnify database before January 2020 (existing samples) (**Fig. 2A; Supplemental Table. S1**). Moreover, to adapt the neural network model to the newly introduced samples, part of which were probably from new biomes, we introduced transfer learning (**Supplemental Fig. S1**) to Meta-Sorter. 34,209 newly introduced microbial samples from 35 biomes (including 8 new biomes)(**Supplemental Table. S1**) and their biome ontology were applied in the transfer learning process to generate the transfer neural network model. During the transfer learning process, the parameters and structures of an existing neural network model could be updated, and the resulting transfer neural network model was suitable for newly introduced samples. Notably, the 34,209 newly introduced samples used here for building the transfer neural network model were those deposited into the MGnify database after January 2020 (newly introduced samples). With the neural network model and the transfer neural network model, Meta-Sorter could decode the samples’ biome labels which were annotated as “Mixed biome” into detailed biome labels (**Fig. 2B**). Besides, Meta-sorter could refine the biome labels to obtain more valuable information for reference (**Fig. 2C**), and correct the mis-annotated samples’ labels to avoid “cascading accumulation” (**Fig. 2D**), as well as other applications, such as classifying the actual sources of ancient DNA (**Fig. 2E**).

### The neural network model decoded the biome information for un-annotated samples accurately

#### The neural network model worked well on different layers of the biome ontology

To disentangle the samples annotated as “Mixed biome”, we need a prediction model that covers as many biomes as possible. Here, we chose 118,592 samples (**Supplemental Table. S1**), which included 134 biomes, and were deposited into the MGnify database before January 2020 (existing samples) to generate a neural network model. The neural network model’s benchmark revealed that the average AUROC, AUPR, and F-max are 0.89, 0.76, and 0.73 respectively. Though the average AUROC on each layer of the biome ontology decreased slightly as the layer increased, the prediction accuracy on each layer exceeded 0.99, indicating the robustness of the neural network model (**Fig. 3A**). Therefore, the neural network model worked well on classifying the existing samples annotated in detail and covered a comprehensive set of biomes, making it feasible to decode the biome labels for samples without detailed biome information.

**Figure 3.**
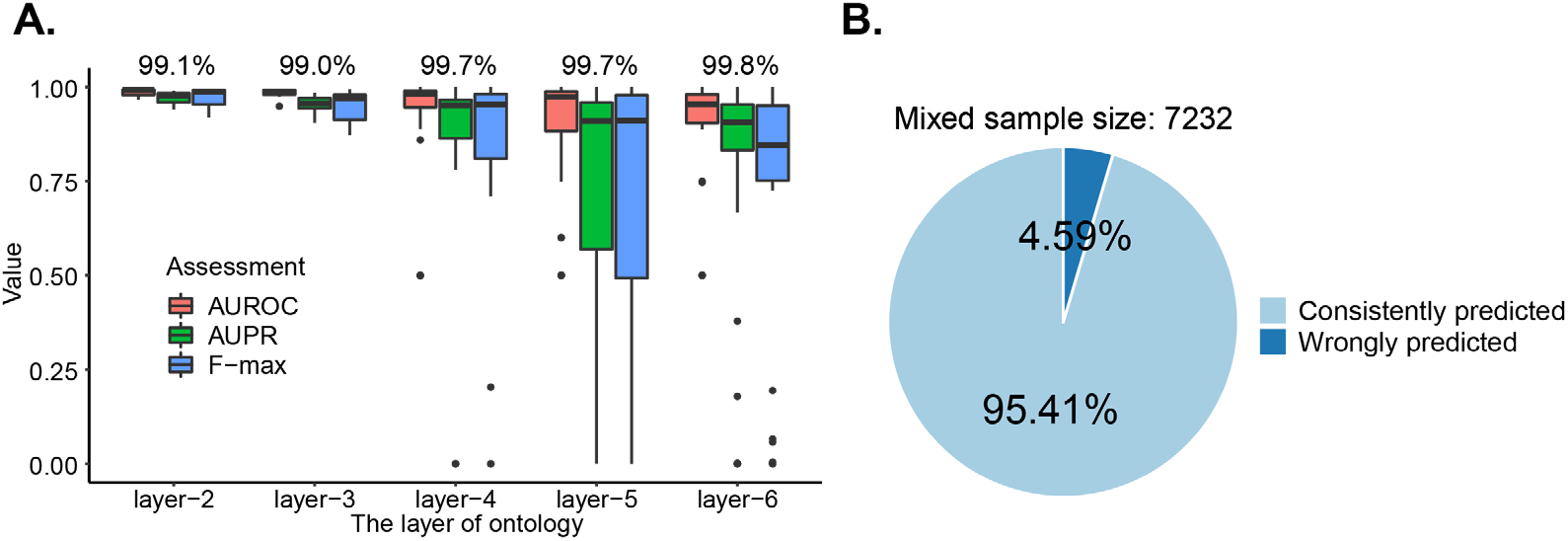
The benchmark of neural network model and the accuracy on decoding the “Mixed biome” labels. (A) 118,592 existing samples were randomly divided into training subset (80%, 94,874 samples) and testing subset (20%, 23,718 samples). The source biome annotation for samples of testing subset were predicted by the neural network model. The boxplots represent AUROC, AUPR and F-max of the neural network model for source biome annotation, categorized by different layers of the biome ontology. The percentages above the boxplots represent the prediction accuracies for each layer of the biome ontology. (B) 7,941 existing samples annotated as “Mixed biomes” were predicted by the neural network model of Meta-Sorter, of which 7,232 samples had reference information in the original literature. Results were based on manually comparing the predicted labels with reference information in the original literature, marked as consistent predicted if the predicted labels were consistent with reference information and marked as wrongly predicted if not consistent.

#### Meta-sorter based on the neural network model decoded the samples’ biome labels annotated as “Mixed biome” into detailed biome labels

We assessed how Meta-Sorter performed on predicting the detailed source biome for samples annotated as “Mixed biome”. We examined 7,941 existing samples from 25 studies annotated as “Mixed biome”, and utilized the neural network model to predict their actual sources (manually curated documented source biome labels as reported in the literature) (**Supplemental Table S2 and S3**). By comparing with the actual sources of these samples, we found that high proportion of biomes predicted by the neural network model were consistent with the actual sources: up to 95.41% of the prediction results of the chosen samples were in line with the original reference information (**Fig. 3B**). Moreover, due to the fact that the neural network model covered comprehensive biomes, it can be applied to disentangle the biome information for samples of a wide range.

#### Case studies on decoded samples from “Mixed biome”

We then focused our examination and in-depth analysis on several representative sets of samples. In the study titled “Home chemical and microbial transitions across urbanization” (study accession: MGYS00005612) (McCall et al. 2020), the researchers collected microbial samples from different rooms in four human environments with different urbanization levels and recorded metadata to explore the role of microbiome structure in urbanization and migration. Despite the high quality and research value, uploaded samples were classified as “Mixed biome”, causing inconvenient access of these samples or even study failure.

We used Meta-Sorter based on the neural network model to disentangle the detailed biome for samples in this built-environment study. By comparing the prediction results (**Supplemental Table S4**) with their actual biomes (Ruiz-Calderon et al. 2016; McCall et al. 2020), we found that despite the presence of non-negligible differences, the categories of the sample sources were substantially consistent (**Fig. 4A, B**), which showed the prediction results by Meta-Sorter were rational and included interesting details listed below.

**Figure 4.**
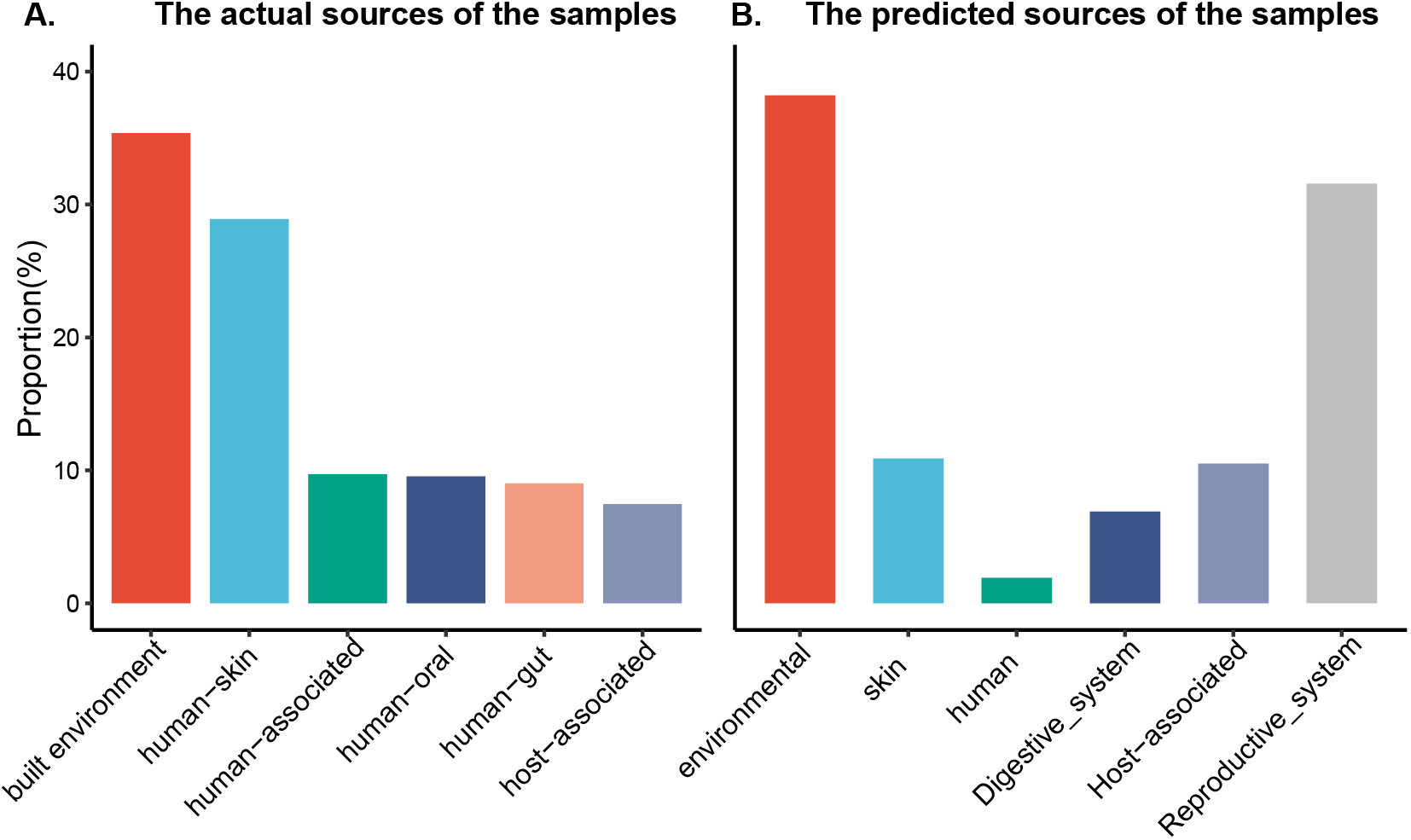
Comparison of predicted sources and actual sources for samples in the case study on microbial communities in built environment for decoding samples from “Mixed biome”. (A) The composition of actual sources and their proportion (%): build environment (35.37), human skin (28.91), human-associated (9.71), human oral (9.54), host-associated (7.46). (B) The composition of predicted sources and their proportion (%): environmental (38.21), human (1.91), skin (10.89), host-associated (10.51), digestive system (6.91), reproductive system (31.56).

Firstly, the original descriptions weren’t refined and accurate enough, while Meta-Sorter got more detailed and concrete results. “Root: Environmental: Aquatic” had the highest percentage of predicted results, but the original description of these samples was “indoor genome,” which belonged to “root: Environmental”. Also noticing this problem, the researchers in this study used a Bayesian approach called SourceTracker (Knights et al. 2011) to estimate the source environments for each group of samples, with results showing that water source, such as domestic water, was the source of a large proportion of the microorganisms, which is highly consistent with Meta-sorter’s results and further validates our model.

Secondly and more importantly, Meta-Sorter could assign samples to biome labels intelligently and automatically, with the biomed labels predicted by Meta-Sorter not included in the manually pre-defined set of biome labels used by SourceTracker. For example, the original description of the samples did not include “reproductive system” (Ruiz-Calderon et al. 2016), whereas in our results, samples sourced from “root: Host-associated: Human: Reproductive system” was as many as 31.56%. The results were highly reasonable, as it was reported in the original literature that some of the samples were collected from floors and walls of bathrooms (using sterile swabs) (Ruiz-Calderon et al. 2016), partly justifying the presence of microbes from the human reproductive system.

### Transfer learning enabled the decoding of the biome information for newly introduced un-annotated samples

#### The limitation of application of the neural network model on newly introduced microbial samples

Though the neural network could perform exceptionally well on disentangling the biome information for un-annotated samples, it should be admitted that all these un-annotated samples were already in the database (existing samples), and their detailed biome information were already known by the neural network model. As the microbial samples in the database accumulated, it would be intriguing if Meta-Sorter could be used to disentangle the biome information for these newly introduced samples. However, the neural network model was only aware of the existing biome information, which was insufficient for such purpose.

We observed that numerous microbial samples were newly introduced into MGnify after January 2020 (newly introduced samples) (**Fig. 1**), we then examined if the neural network model of Meta-Sorter could be utilized to predict the actual sources for these samples. Here, we have collected 32 newly introduced studies, which included 34,209 samples annotated with detailed information, and 10,862 samples from 10 studies annotated as “Mixed biome”. Meta-Sorter based on the neural network model has been applied on the 34,209 samples annotated with detailed information, and compared with the independent neural network model which was constructed solely on these 34,209 samples. The average AUROC of the neural network model is 0.872 while the independent neural network model is 0.989 (p_Wilcox_=2.22×10^−16^; **Fig. 5A**), a pivotal reason for which was that the distribution between existing samples and newly introduced samples had significant differences (**Fig. 5B**). This phenomenon restricted the applicability of existing neural network model on the newly introduced samples.

**Figure 5.**
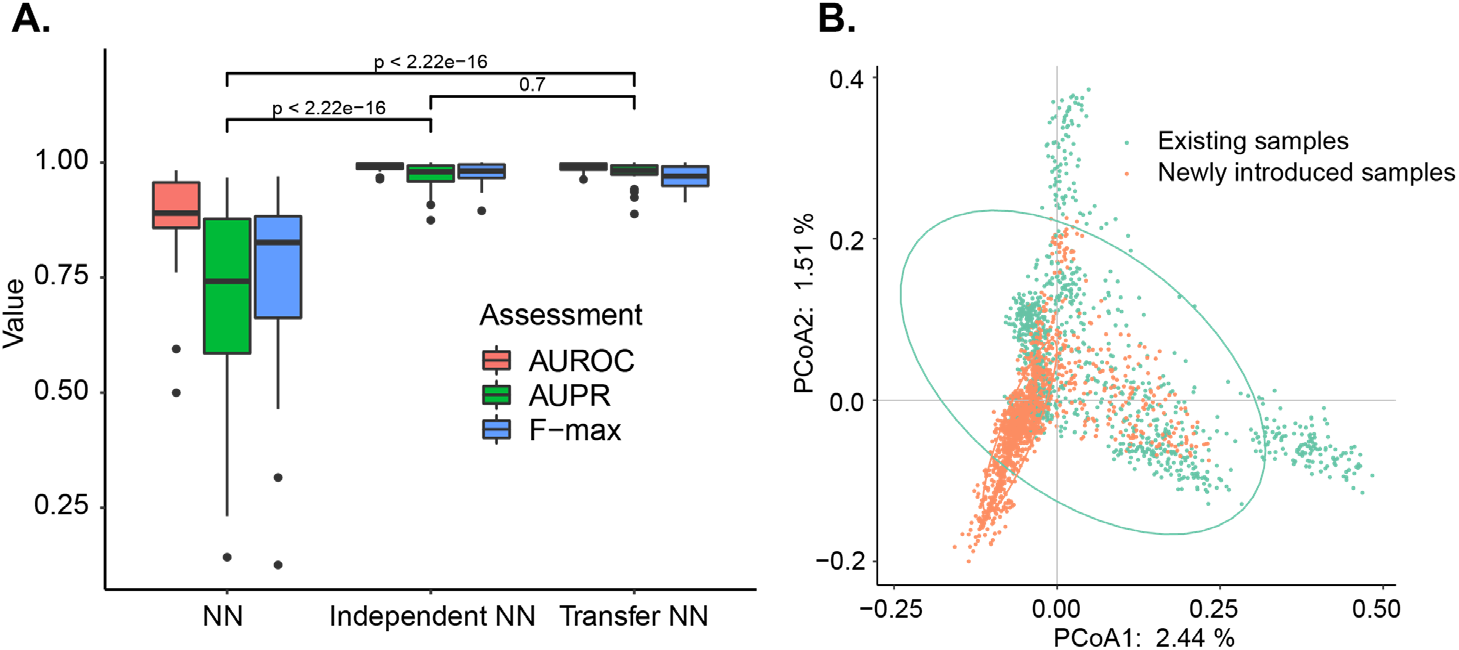
The benchmark of different models and the heterogeneity of existing samples and newly introduced samples. (A) Comparison of different models in source biome annotation for newly introduced samples. 34,209 newly introduced samples with detailed biome information were randomly divided into training subset (80%, 27,355 samples) and testing subset (20%, 6,854 samples), the samples of testing subset were predicted by NN model, independent NN model and transfer NN model respectively. The boxplots represent AUROC, AUPR and F-max of the neural network model for source biome annotation among three models. *P* < 0.05; **, P < 0.01; ***, *P* < 0.005; Wilcoxon signed-rank test. (B) The different distribution of existing samples and newly introduced samples. The confidence level is 95%. NN, neural network.

However, as more and more samples annotated as “Mixed biome” would be introduced into the public database like MGnify (**Fig. 1**), it’s desirable to find out solution for exceeding the effect of the heterogeneity of datasets and disentangling the biome information of those samples in “Mixed biome”. To solve this problem, we introduced transfer learning (Cai et al. 2020) to update the neural network model and generated a transfer neural network model, which could be utilized for disentangling the biome information for these newly introduced microbial samples which were annotated as “Mixed biome” (**Fig. 2A; Supplemental Fig. S1**). The average AUROC of transfer neural network model was 0.989, outperformed than the neural network model (p_Wilcox_=2.22×10^−16^) and as good as the independent neural network model (p_Wilcox_=0.7), with the consistent results of AUPR and F-max (**Fig. 5A**). This result indicated that transfer learning could efficiently exceed the limitation for application of neural network caused by heterogeneity between the existing samples and the newly introduced samples.

#### Transfer learning enhanced the adaptability of the neural network model to newly introduced biome

In addition to the newly introduced samples which only include the existing biomes, there are a considerable number of newly introduced biomes, despite the obvious fact that the neural network model was not suitable for classifying these samples from new biomes. The first and foremost reason is that remodeling in each newly introduced biome is unrealistic due to the enormous computing resources and time required. There’s another reason that, if the amount of data newly added to the database in a certain year is rather small, or when the researchers aim to excavate specialized data (e.g., marine microbial data mining), the independent model cannot be adequately trained due to the limited size of dataset. In these contexts, transfer learning scheme is suitable since less time and computing resources are required to generate a reliable transfer neural network model. Therefore, we applied transfer learning to adapt the neural network model to 34,209 newly introduced samples from 35 biomes (including 3,083 samples from 8 newly introduced biomes) to generate the transfer neural network model (**Fig. 6A**).

**Figure 6.**
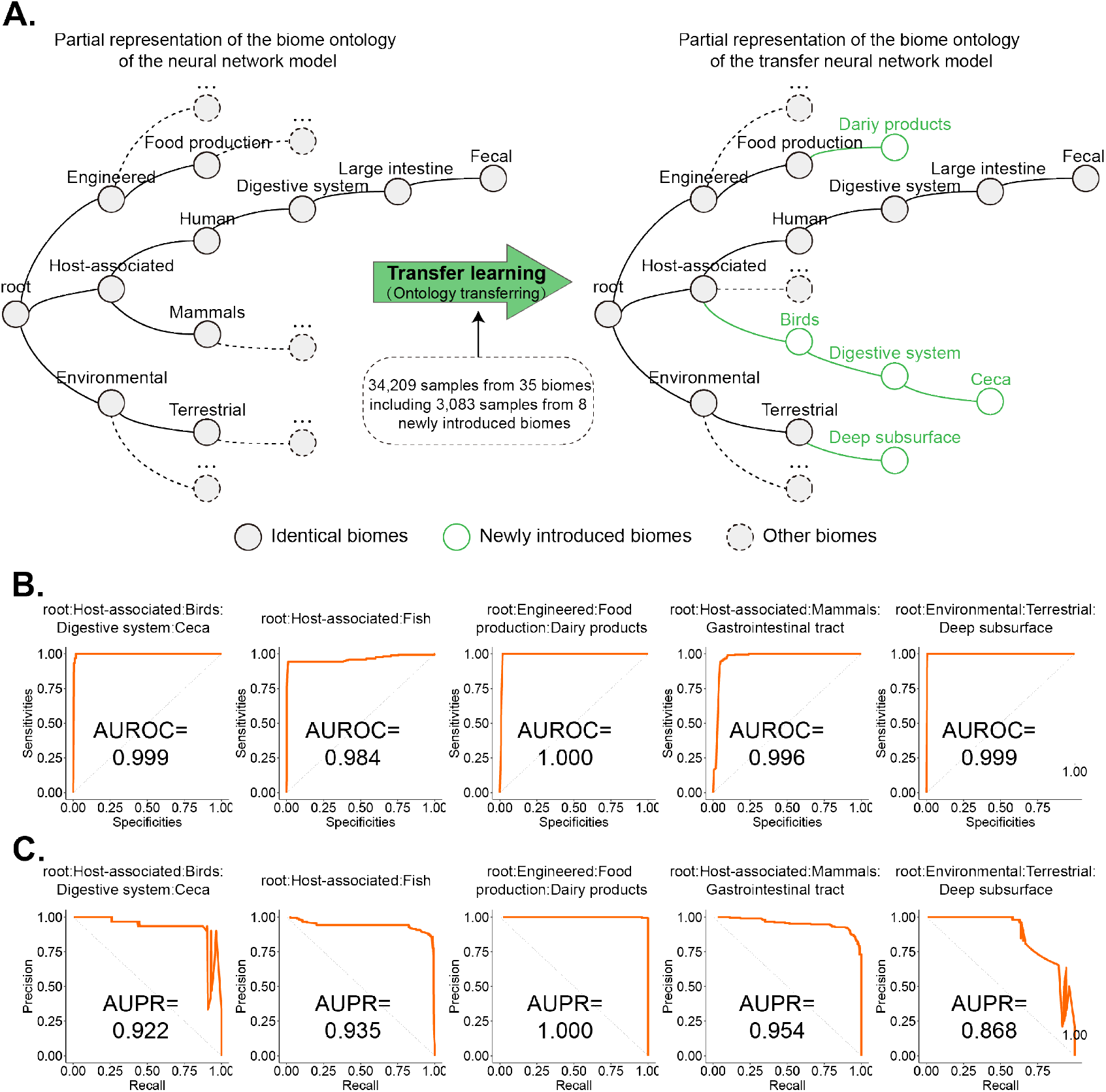
The robust adaptation of Meta-Sorter to the newly introduced samples. (A) The left panel shows that the partial representation of the biome ontology of existing samples used to constructed the neural network model, and the right panel shows that the partial representation of the biome ontology of the newly introduced samples used for transfer learning to generate the transfer neural network model. The existing samples included 118,592 samples from 134 biomes and the newly introduced samples included 34,209 samples from 35 biomes (including 8 newly introduced biomes, e.g. birds related biomes). (B, C) The AUROC and AUPR assessments of Meta-Sorter on representative newly introduced biomes.

We assessed the AUROC and AUPR of transfer neural network model predicting the newly introduced biomes, and found that the transfer neural network model worked well on those biomes (**Fig. 6B, C**). There are plenty of cases that support the superiority of transfer neural network. For example, we noticed that there was a new biome out of the 35 biomes which was annotated as “root: Host-associated: Birds: Digestive system: Ceca”. However, only “root: Host-associated” was introduced in the neural network model, which obviously was not suitable for those newly introduced biomes, while by implementing transfer learning to the neural network model with samples in the new biome, the transfer neural network model had high prediction accuracy (AUROC = 0.999 and AUPR = 0.868) on the biome. In short, transfer learning enhanced the adaptability of the neural network model to the newly introduced biomes, the robustness of the model has been improved.

#### Prediction of the detailed source biome for samples annotated as “Mixed biome” from newly introduced studies

Apart from the samples with detailed biome labels, there were a proportion of newly introduced samples without biome information and annotated as “Mixed biome”. Due to the existence of differences among existing samples and newly introduced samples, the neural network model may not be appropriate for decoding their biome labels. Therefore, we assessed the accuracy of Meta-Sorter based on the transfer neural network model on predicting the detailed source biome for newly introduced samples annotated as “Mixed biome”. We chose 10,862 newly introduced samples from 10 studies annotated as “Mixed biome” (**Supplemental Table. S2**), and utilized the transfer neural network model to disentangle the detailed biomes for these samples, then compared with their actual sources from the original literature (**Supplemental Table S5**). We found that 97.62% of these samples were assigned to detailed biomes which were consistent with their actual sources (**Fig. 7A**), indicating that the transfer neural network model could effectively disentangle the biome information for those samples without detailed annotations.

**Figure 7.**
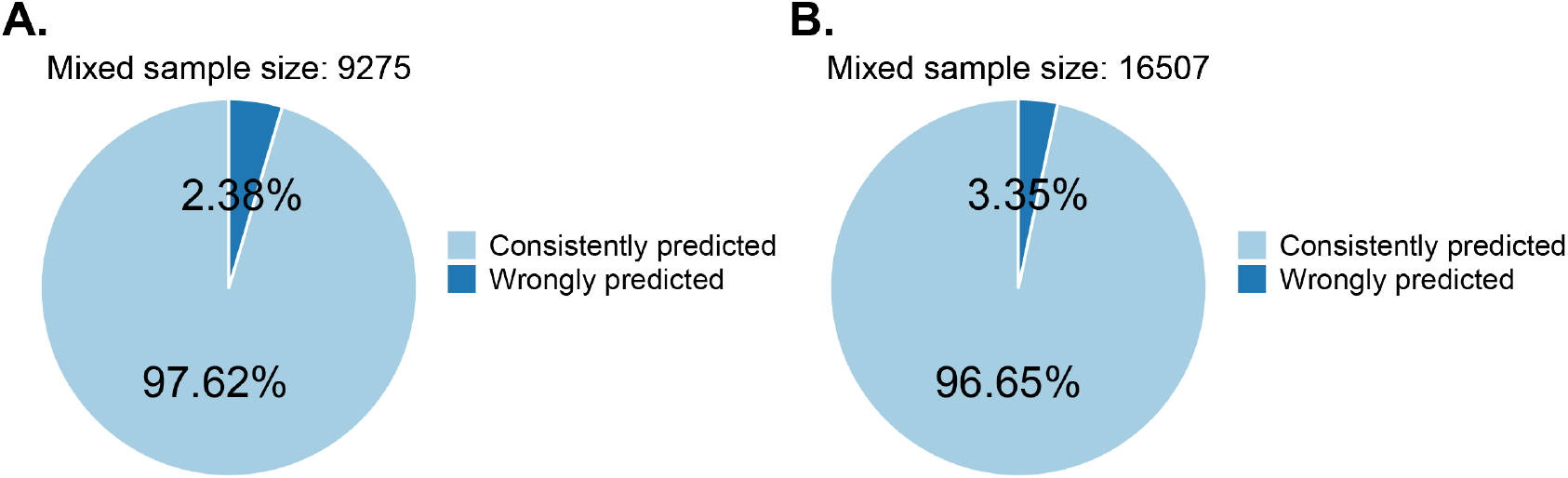
The accuracy of Meta-Sorter on decoding samples annotated as “Mixed biome”. (A) 10,862 newly introduced samples annotated as “Mixed biomes” were predicted by the transfer neural network model of Meta-Sorter, of which 9,275 samples had reference information in the original literature. We manually compared the predicted labels with reference information in the original literature, and marked as consistent predicted if the predicted labels were consistent with reference information, while marked as wrongly predicted if not consistent. (B) The general prediction accuracy of Meta-Sorter on the samples in “Mixed biome”, including those from both existing samples and newly introduced samples. Meta-Sorter altogether predicted 18,803 samples annotated as “Mixed biome” based on the neural network model and the transfer neural network model, of which 16,507 samples had reference information in the original literature.

Combined the results on existing samples and newly introduced samples, we found that Meta-Sorter based on neural network model and transfer neural network could largely solving the missing biome labeling problem: 16,507 samples out of 18,803 total samples had reference information in the original literature, of which 96.65% (15,954 samples) could be consistently predicted by Meta-Sorter (**Fig. 7B**). This demonstrated the effectiveness of Meta-Sorter.

### Meta-Sorter could refine the biome annotation for under-annotated and mis-annotated samples

In addition to decoding samples labeled as “Mixed biome”, Meta-Sorter can also classify data too high in the classification hierarchy to be refined. In study “anchialine metagenome raw sequence reads” (MGYS00005510), researchers collected benthic and water column samples from nine anchialine habitats in the Hawaiian Archipelago and identified environmental factors driving microbial diversity. Despite the arduous sample-collecting procedure and genomes information extremely exploitable for microbiomics and metagenomics researchers, all samples were labeled as “Root: Environmental: Aquatic,” and even the sample descriptions lacked precise site information. We utilized Meta-Sorter based on transfer learning to refine the label of 250 samples (**Supplemental Table. S6**) in this study with results showing as follows:

Firstly, Meta-Sorter successfully deciphered the sample information for more refined classification labels. 85.6% of the predicted results from Meta-Sorter reached to the classification of layer5 or layer6 (**Fig. 8A**), while the original classification label for all samples was “root: Environmental: Aquatic”, which may provide added offer additional information to other researchers when mining data at finer classification levels.

**Figure 8.**
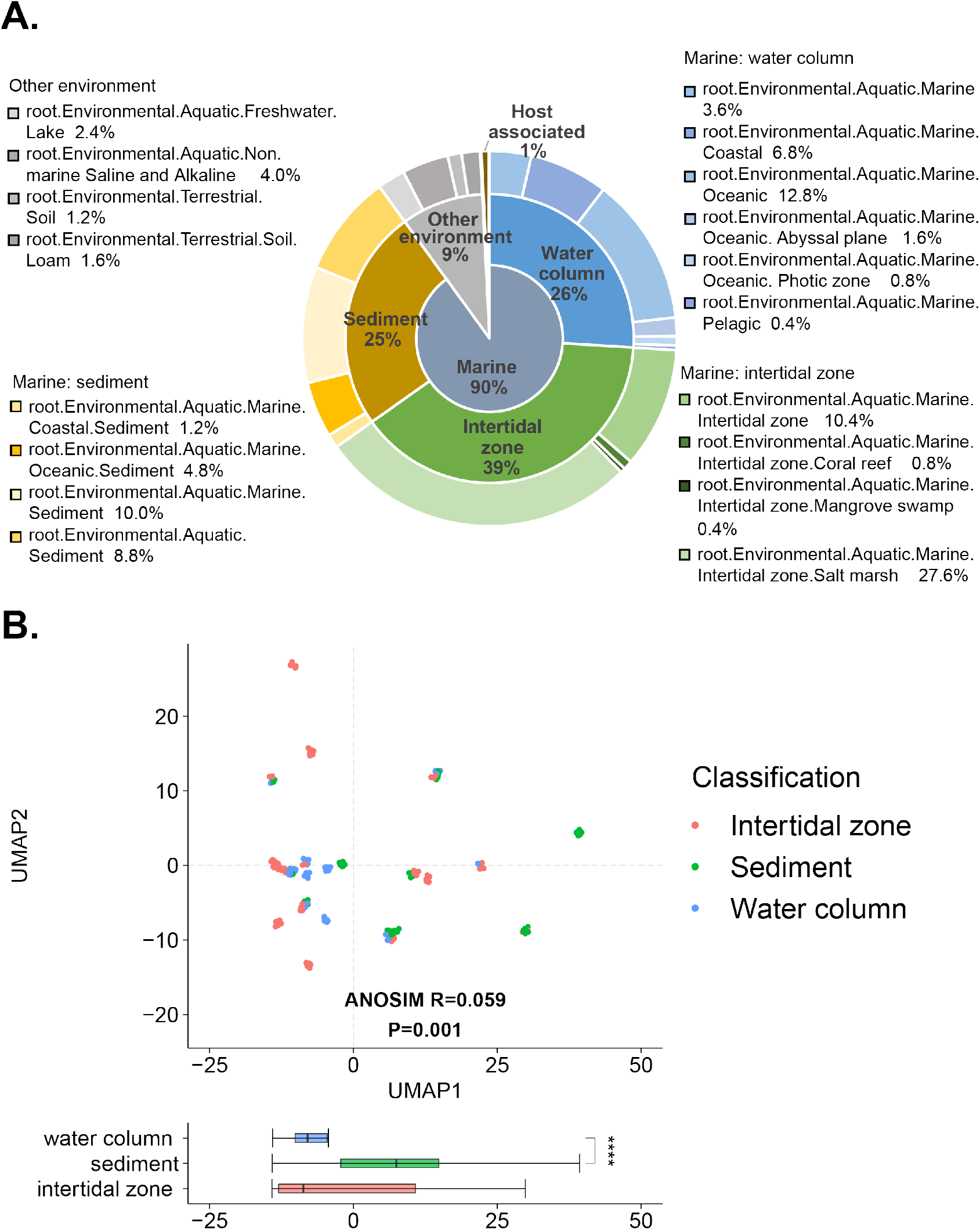
Meta-Sorter refined the source labels of representative samples from a case study on marine samples. (A) The detailed biome contribution predicted by Meta-Sorter. The results were manually compartmentalized into three sources: water column, intertidal zone and sediment, more detailed sources and proportions were as well as shown under the three sources. (B) The distribution of the samples with the predicted three-category source labels (water column, sediment and intertidal zone) as the samples’ actual sources (ANOSIM, *R = 0.059, P = 0.001*). The box-plot showed the distribution of samples on the x-axis, demonstrating substantial differences between water column and sediment sample (Wilcoxon, *p < 0.0001*).

Secondly, Meta-Sorter’s results were rational and indicated extra clues for this study. The sampling environment is anchialine open pools or ponds and caves, which are part of the marine/anchialine ecosystem, whereas Meta-Sorter predicted that 90% of the samples would be classified as “root: Environmental: Aquatic: Marine.” When studying the variations in community structure in distinctive environments, the researchers divided the samples into two categories: water column and benthic, with Meta-Sorter predicting that 26% of the samples correspond to the “water column” portion of the original labels, and 25% of the samples correspond to the “benthic” section of the original labels (**Fig. 8A**). Collectively, the aforementioned results demonstrate the validity of our predictions.

Interestingly, Meta-Sorter’s results show clues why water columns and benthos shared similar communities. A significant proportion of predictions (39%) belong to “root: Environmental: Aquatic: Marine: Intertidal zone”, which denotes the area above water level at low tide and underwater at high tide, where environmental and biome variability is high, including the biomes of neritic, deep zones, and seabed (Menge 2000; Levin et al. 2001). In the NMDS analysis, the researchers found higher numbers of shared communities and limited variation in the structure of the water column and benthic communities at the same sampling site, most likely due to the fact that both taxa contained a substantial proportion of samples from the “intertidal zone” (**Fig. 8B**). If the researcher excludes that part of the sample or reanalyzes it by the three categories, additional creative conclusions may be drawn.

Finally, predictions that appear inconsistent with the sampling environment may indicate biome migration. Of the Meta-Sorter predictions, 1% of the samples were identified as host associated, and 9% were classified as environments other than marine, such as “Freshwater Lake” and “Non-marine Saline and Alkaline”, which can be partially attributed to the sampling sites. Furthermore, the biomes predicted as the label of host associated and other environments may have some tendency to have inhabited both ecological niches, indicating community migration between diverse environments and hosts (Fuhrman et al. 2015). In addition, there is a non-negligible possibility that this inconsistency is due to human factors such as contamination of the samples resulting in labeling errors, which also reveals Meta-sorter’s potential to resolve the mis-annotation problems.

Collectively, Meta-Sorter could not only disentangle the detailed biome information for un-annotated samples, but also refine the biome annotation for under-annotated and mis-annotated samples. More interestingly, Meta-Sorter was able to assign samples to biome labels in a highly intelligent and automatic manner, telling us more than those reported in the original literature.

## Discussion

It has become an increasing difficult problem for data mining from publicly available microbiome samples, largely due to three kinds of inaccurately annotated samples (un-annotated samples, under-annotated samples and mis-annotated samples), which has led to the inability for biomarker discovery or even “cascading accumulation” of errors. In this study, we have designed Meta-Sorter, a neural network and transfer learning enabled AI method for improving the biome labeling of thousands of microbial community samples with inaccurate biome information. A comprehensive neural network model was established, and the transfer neural network model based on transfer learning was then introduced to decode information for newly introduced biome, both of which can effectively solve the problems caused by the mentioned three kinds of samples. Results have shown that out of 16,507 samples that have no detailed biome annotations, 15,954 (96.65%) could be consistently predicted, largely solving the biome labeling problem for inaccurately annotated samples. Interestingly, Meta-Sorter was able to assign samples to biome labels in a highly intelligent and automatic manner, telling us more about microbial community samples than those reported in the original literature. Moreover, Meta-Sorter has a broad spectrum of applications, such as indicating the potential of microbial community migration, disentangling the hidden information of ancient microbial community samples and examining whether a sample is contaminated.

Meta-Sorter could be further improved when more microbial community samples and their meta-data are accumulated. With the accumulation of samples from more diverse biomes, Meta-Sorter’s model could include more biome information for more accurate biome labeling. Additionally, for under-annotated samples and mis-annotated samples, mining Meta-Sorter’s results might reveal intricate but important connections among samples. Finally, the idea of Meta-Sorter using both neural network and transfer learning for sample annotation might be expanded to other domains of biological data mining in a wide range of contexts such as gene mining.

To conclude, we have improved the completeness of biome label information for tens of thousands of microbial community samples, solving the problem of errors’ “cascading accumulation”, facilitating a wide range of applications, including sample classification, source tracking and novel knowledge discovery, from millions of microbiome samples.

## Methods

### Datasets

We examined 118,592 samples from 1,447 studies introduced into MGnify before January 2020 to generate the neural network model, 25 studies annotated as “Mixed biome” before January 2020 which included 7,941 samples. 34,209 samples newly introduced into MGnify after January 2020, which belonged to 32 studies, were applied for implementing transfer learning to the neural network model to generate the transfer neural network model, and we also introduced 10 studies annotated as “Mixed biome” after January 2020 which included 10,862 samples

### Process of model construction and transfer learning

Meta-Sorter is based on the ontology-aware neural network model, combine with transfer learning.

Data processing process: First import the microbial data (bacteria taxonomy abundance table), establish a mapping relationship between the abundance table and the phylogenetic tree, construct a regular abundance matrix, and then standardize the abundance matrix and convert it into relative abundance matrix. Then the relative abundance matrix is standardized by Z-score to become a standard abundance matrix. Ontology-aware neural network modeling process: The neural network model is built based on mimicking the biome ontology structure: each node represents a biome, and the layers from bottom to top represents the biome ontology structure (for example, the biome “root-Host-associated-Human-Digestive_system-Large_intestine-Fecal” is represented in the model as: layer-1 represents “root”, layer-2 represents “Host-associated”, …, layer-6 represents “Fecal”). The neural network thus been built is referred to as the general neural network model (general model in short).

Transfer learning process: The existing model could be divided into bottom-level nodes and top-level nodes. The bottom-level nodes have the potential to be applied in emerging datasets, while the top-level nodes can only be applied to the existing datasets. Firstly, lock the bottom nodes so that they do not participate in the transferring process, encode and introduce the new community structure, and change the structure and weights between the top-level nodes. This process is transfer process (Transfer). Input the microbial data from the new community, through the forward algorithm and the backward algorithm, iteratively update the top-level parameters until convergence, then the top-level nodes can be applied to the new datasets, this process is fast adaptation process (Fast adaptation). Unlock the bottom-level nodes, use new microbial data and the training of forward and backward algorithms, and iteratively update the parameters of the bottom-level nodes until convergence. This process is fine-tuning process (Fine-tuning).

### Performance measures

For AUROC, AUPR and F-max computation, we set the threshold ∈[0,1] with a step size of 0.01, the result of logical operation is 1 if the contribution of node is greater than the threshold, else 0. We calculated True Positive, True Negative, False Positive, False Negative, for calculating True Positive Rate and False Positive Rate, Precision, Recall and F1-measure at ever threshold, then we obtain the AUC curve and PR curve. By calculating area under the AUC curve as AUROC and area under the PR curve as AUPR of each node, F-max stands for the maximal F1-measure. Each node represents an ecological classification of a community.

### Assessments of Meta-Sorter

We accessed the neural network model by applying five-fold cross-validation to the 118,592 samples collected from 134 biomes, the neural network model with the best performance was used for Meta-Sorter.

We accessed the transfer neural network and the independent neural network model by applying five-fold cross-validation (80% samples as source to construct the independent neural network model and to implement transfer learning to the neural network model for generating the transfer neural network model, the rest 20% samples were used to assess performance of the models) to the 34,209 samples from 32 studies, the transfer neural network model with the best performance was used for Meta-Sorter.

## Data Access

The datasets analysed during the current study are publicly available in EBI-MGnify (https://www.ebi.ac.uk/metagenomics). The accession numbers are included in Supplemental Table. S1.

## Acknowledgments

This work was partially supported by National Natural Science Foundation of China grant 32071465, 31871334, and 31671374, and the China Ministry of Science and Technology’s National Key R&D Program grant (No. 2018YFC0910502). We thank Hui Chong for the technical assistance in transfer learning process.

## Author contribution

NW and KN conceived the idea; NW and TW performed the study; NW and TW prepared and analyzed the data; NW and TW wrote the manuscript. All authors read and approved the final manuscript.

## References

Bahram M, Hildebrand F, Forslund SK, Anderson JL, Soudzilovskaia NA, Bodegom PM, Bengtsson-Palme J, Anslan S, Coelho LP, Harend H et al. 2018. Structure and function of the global topsoil microbiome. Nature 560: 233–237.

Bolyen E Rideout JR Dillon MR Bokulich NA Abnet CC Al-Ghalith GA Alexander H Alm EJ Arumugam M Asnicar F et al. 2019. Reproducible, interactive, scalable and extensible microbiome data science using QIIME 2. Nat Biotechnol 37: 852–857.

Brooks B, Firek BA, Miller CS, Sharon I, Thomas BC, Baker R, Morowitz MJ, Banfield JF. 2014. Microbes in the neonatal intensive care unit resemble those found in the gut of premature infants. Microbiome 2: 1.

Cai C, Wang S, Xu Y, Zhang W, Tang K, Ouyang Q, Lai L, Pei J. 2020. Transfer Learning for Drug Discovery. J Med Chem 63: 8683–8694.

Escobar-Zepeda A, Vera-Ponce de León A, Sanchez-Flores A. 2015. The Road to Metagenomics: From Microbiology to DNA Sequencing Technologies and Bioinformatics. Front Genet 6 : 348.

Fuhrman JA, Cram JA, Needham DM. 2015. Marine microbial community dynamics and their ecological interpretation. Nat Rev Microbiol 13: 133–146.

Gaulke CA, Sharpton TJ. 2018. The influence of ethnicity and geography on human gut microbiome composition. Nat Med 24: 1495–1496.

Gonzalez A, Navas-Molina JA, Kosciolek T, McDonald D, Vázquez-Baeza Y, Ackermann G, DeReus J, Janssen S, Swafford AD, Orchanian SB et al. 2018. Qiita: rapid, web-enabled microbiome meta-analysis. Nat Methods 15: 796–798.

Halfvarson J, Brislawn CJ, Lamendella R, Vázquez-Baeza Y, Walters WA, Bramer LM, D’Amato M, Bonfiglio F, McDonald D, Gonzalez A et al. 2017. Dynamics of the human gut microbiome in inflammatory bowel disease. Nat Microbiol 2: 17004.

Keegan KP, Glass EM, Meyer F. 2016. MG-RAST, a Metagenomics Service for Analysis of Microbial Community Structure and Function. Methods Mol Biol 1399: 207–33.

Kelley ST, Gilbert JA. 2013. Studying the microbiology of the indoor environment. Genome Biol 14: 202.

Knights D, Kuczynski J, Charlson ES, Zaneveld J, Mozer MC, Collman RG, Bushman FD, Knight R, Kelley ST. 2011. Bayesian community-wide culture-independent microbial source tracking. Nat Methods 8: 761–763.

Kriegeskorte N, Golan T. 2019. Neural network models and deep learning. Curr. Biol. 29: 231–236.

Lax S, Sangwan N, Smith D, Larsen P, Handley KM, Richardson M, Guyton K, Krezalek M, Shogan BD, Defazio J et al. 2017. Bacterial colonization and succession in a newly opened hospital. Sci Transl Med 9: 391.

Levin LA, Boesch DF, Covich A, Dahm C, Erséus C, Ewel KC, Kneib RT, Moldenke A, Palmer MA, Snelgrove P et al. 2001. The Function of Marine Critical Transition Zones and the Importance of Sediment Biodiversity. Ecosystems 4: 430–451.

Lu J, Salzberg SL. 2020. Ultrafast and accurate 16S rRNA microbial community analysis using Kraken 2. Microbiome 8: 124.

McCall L-I, Callewaert C, Zhu Q, Song SJ, Bouslimani A, Minich JJ, Ernst M, Ruiz-Calderon JF, Cavallin H, Pereira HS et al. 2020. Home chemical and microbial transitions across urbanization. Nat Microbiol 5: 108–115.

Menge BA. 2000. Top-down and bottom-up community regulation in marine rocky intertidal habitats. J Exp Mar Biol Ecol 250: 257–289.

Mitchell AL, Almeida A, Beracochea M, Boland M, Burgin J, Cochrane G, Crusoe MR, Kale V, Potter SC, Richardson LJ et al. 2020. MGnify: the microbiome analysis resource in 2020. Nucleic Acids Res 48: 570–578.

Ruiz-Calderon JF, Cavallin H, Song SJ, Novoselac A, Pericchi LR, Hernandez JN, Rios R, Branch OH, Pereira H, Paulino LC et al. 2016. Walls talk: Microbial biogeography of homes spanning urbanization. Sci Adv 2: 2.

Sayers EW, Beck J, Bolton EE, Bourexis D, Brister JR, Canese K, Comeau DC, Funk K, Kim S, Klimke W et al. 2021. Database resources of the National Center for Biotechnology Information. Nucleic Acids Res 49: 10–17.

Wang Z, Han M, Li E, Liu X, Wei H, Yang C, Lu S, Ning K. 2020. Distribution of antibiotic resistance genes in an agriculturally disturbed lake in China: Their links with microbial communities, antibiotics, and water quality. J Hazard Mater 393: 122426.

Wu S, Sun C, Li Y, Wang T, Jia L, Lai S, Yang Y, Luo P, Dai D, Yang Y-Q et al. 2020. GMrepo: a database of curated and consistently annotated human gut metagenomes. Nucleic Acids Res 48: 545–553.

